# The entropy of resting-state neural dynamics is a marker of general cognitive ability in childhood

**DOI:** 10.1101/2023.08.08.552448

**Authors:** Natalia Zdorovtsova, Edward J. Young, Danyal Akarca, Alexander Anwyl-Irvine, The RED Team, The CALM Team, Duncan E. Astle

## Abstract

1

Resting-state network activity has been associated with the emergence of individual differences across childhood development. However, due to the limitations of time-averaged representations of neural activity, little is known about how cognitive and behavioural variability relates to the rapid spatiotemporal dynamics of these networks. Magnetoencephalography (MEG), which records neural activity at a millisecond timescale, can be combined with Hidden Markov Modelling (HMM) to track the spatial and temporal characteristics of transient neural states. We applied HMMs to resting-state MEG data from (n = 46) children aged 8-13, who were also assessed on their cognitive ability and across multiple parent-report measures of behaviour. We found that entropy-related properties of participants’ resting-state time-courses were positively associated with cognitive ability. Additionally, cognitive ability was positively correlated with the probability of transitioning into HMM states involving fronto-parietal and somatomotor activation, and negatively associated with a state distinguished by default-mode network suppression. We discuss how using dynamical measures to characterise rapid, spontaneous patterns of brain activity can shed new light on neurodevelopmental processes implicated in the emergence of cognitive differences in childhood.

**Significance Statement:** There is increasing evidence that the function of resting-state brain networks contributes to individual differences in cognition and behaviour across development. However, the relationship between dynamic, transient patterns of switching between resting-state networks and neurodevelopmental diversity is largely unknown. Here, we show that cognitive ability in childhood is related to the complexity of resting-state brain dynamics. Additionally, we demonstrate that the probability of transitioning into and remaining in certain ‘states’ of brain network activity predicts individual differences in cognitive ability.

## 2 Introduction

### 2.1 The emergence of resting-state networks across childhood development

Brain activity is characterised by intrinsic dynamics in the form of spatially-coherent and spontaneous fluctuations, which form resting-state networks (RSNs). RSNs were first discovered in the form of functionally-correlated features of fMRI timeseries during resting-state scans (Biswal et al., 1995; Lowe et al., 1998; Xiong et al., 1999; Cordes et al., 2000), and later taxonomised across a wide range of neuroimaging modalities (Kiviniemi et al., 2003; van den Heuvel et al., 2009; Allen et al., 2011). Further studies revealed a correspondence between the brain’s functional architecture across tasks and at rest, indicating that these networks support distinct features of sensory processing, cognition, and behaviour (Smith et al. 2009). As such, RSNs are thought to have emerged across evolutionary timescales, and are known to develop from infancy through adolescence, in order to serve specific functions (Yeo et al., 2011). Understanding the nature of spontaneous patterns of brain activity, and how they develop, is therefore an important challenge within developmental cognitive neuroscience.

### 2.2 Resting-state characteristics predict individual differences in cognition and behaviour

Divergent RSN development has been observed in those with neurodevelopmental conditions. Typical childhood development is characterised by a gradual segregation between the default-mode network and task-positive networks across the brain. Functional over-connectivity between these networks is associated with cognitive and behavioural difficulties, particularly Attention-Deficit Hyperactivity Disorder (ADHD) (Cortese et al., 2012; Sripada et al., 2014; Francx et al., 2015; Cai et al., 2018; de Lacy & Calhoun, 2018; Jones et al., 2022). Children with conditions like ADHD have also been found to demonstrate profiles of lower fronto-parietal activation at rest (Cortese et al., 2012; Castellanos & Proal, 2012), as have those diagnosed with intellectual disabilities in childhood (Ma et al., 2021). Given the fact that RSN function is implicated in developmental divergence, there is value in outlining the precise mechanisms by which RSN differences might correspond to population-level variations in cognitive ability and behavioural difficulty.

### 2.3 Using dynamical measures to characterise the spatiotemporal properties of spontaneous brain activity

As discussed previously, spontaneous functional network activity can be described using static, time-averaged representations of neural activation. However, computational models and empirical data suggest that the brain engages in rapid switching between distinct states of functional connectivity (Rabinovich et al., 2008; Nachstedt & Tetzlaff, 2017). In other words, states of neural activity are not merely situated in space, but also in time, and transient, recurrent patterns of activity could represent a meaningful mode by which information is processed by the brain. One limitation of time-averaged representations is that they are unable to describe the transient temporal dynamics of brain activity. An emerging field dedicated to measuring these rapid dynamics strives to overcome some of the interpretability constraints of time-averaged functional network identification techniques by proposing a new set of data-driven, temporally-embedded methods of representing neural activity.

One of these relatively novel methods is Hidden Markov Modelling (HMM), an unsupervised machine learning technique that reconstructs a sequence of patterns as a system of temporally-discrete states. Previously, HMMs have been used to extract the underlying dynamical properties of neural data from MEG (Baker et al., 2014; Vidaurre et al., 2018b; Quinn et al., 2018, Hawkins et al., 2019), EEG (Obermaier et al., 2001; Williams et al., 2018; Dash & Kolekar, 2020; Marzetti, 2023), and fMRI (Duan et al., 2005; Dang et al., 2017; Goucher-Lambert & McComb, 2019; Hussain et al., 2023) at rest and in task settings. To our knowledge, there have not been any studies of how resting-state neural dynamics vary in a developmental sample, or how these dynamics relate to the emergence of diverse profiles of behaviour and cognitive ability.

### 2.4 The current study

In the current study, we used resting-state MEG data from children aged 8-13 to test how the temporal properties of transient neural dynamics vary with age, gender, behavioural difficulties, and general cognitive ability. We also explored the extent to which the complexity of individual participants’ resting-state time-courses varied across these neurodevelopmental features of interest.

## 3 Materials and Methods

### 3.1 Participants

Our MEG analysis sample, following exclusions (n = 46), included children aged 8-13 (M = 10.09, SD = 1.19). Children were all part of ongoing studies at the MRC Cognition and Brain Sciences Unit, and all underwent an identical MEG protocol. Specifically, all the children were originally part of two studies, either the Resilience in Education and Development (RED) study (n = 40, PRE.2017.102) or the Centre for Attention, Learning, and Memory (CALM) cohort (n = 6, 22/WM/0082), and agreed to take part in this MEG session. Combined, this sample reflects a range of common behavioural difficulties typically seen in mainstream education.

### 3.2 Cognitive Assessments and Behavioural Questionnaires

#### 3.2.1 Wechsler Abbreviated Scales of Intelligence II – Matrix Reasoning Subtest (WASI-II MR)

The Matrix Reasoning subtest of the Wechsler Abbreviated Scales of Intelligence II is a general measure of cognitive ability and executive function. In this subtest, children are presented with incomplete matrices of images and asked to select an image that would suitably complete each matrix from a choice of four options. Children aged 9 years and older complete a possible total of 30 matrices, which become progressively more difficult. The matrix reasoning test is finished when the child produces three incorrect answers in a row. Trials correct were converted to age-standardised T-scores.

#### 3.2.2 Strengths and Difficulties Questionnaire

The Strengths and Difficulties Questionnaire (SDQ) asked parents/carers to answer 25 questions measuring a variety of behavioural challenges (with possible responses being ‘Not True,’ ‘Somewhat True,’ and ‘Certainly True’) based on their child’s behaviour in the six months prior to assessment. A total SDQ score is calculated, in addition to scores for five behavioural subscales: Hyperactivity, Conduct Problems, Emotional Regulation Problems, Peer Relationship Problems, and Prosocial Behaviour (see Table 1). See Supplementary Figure 1 for plots representing how cognitive, behavioural, and demographic traits were distributed across our combined sample.

**Figure 1:**
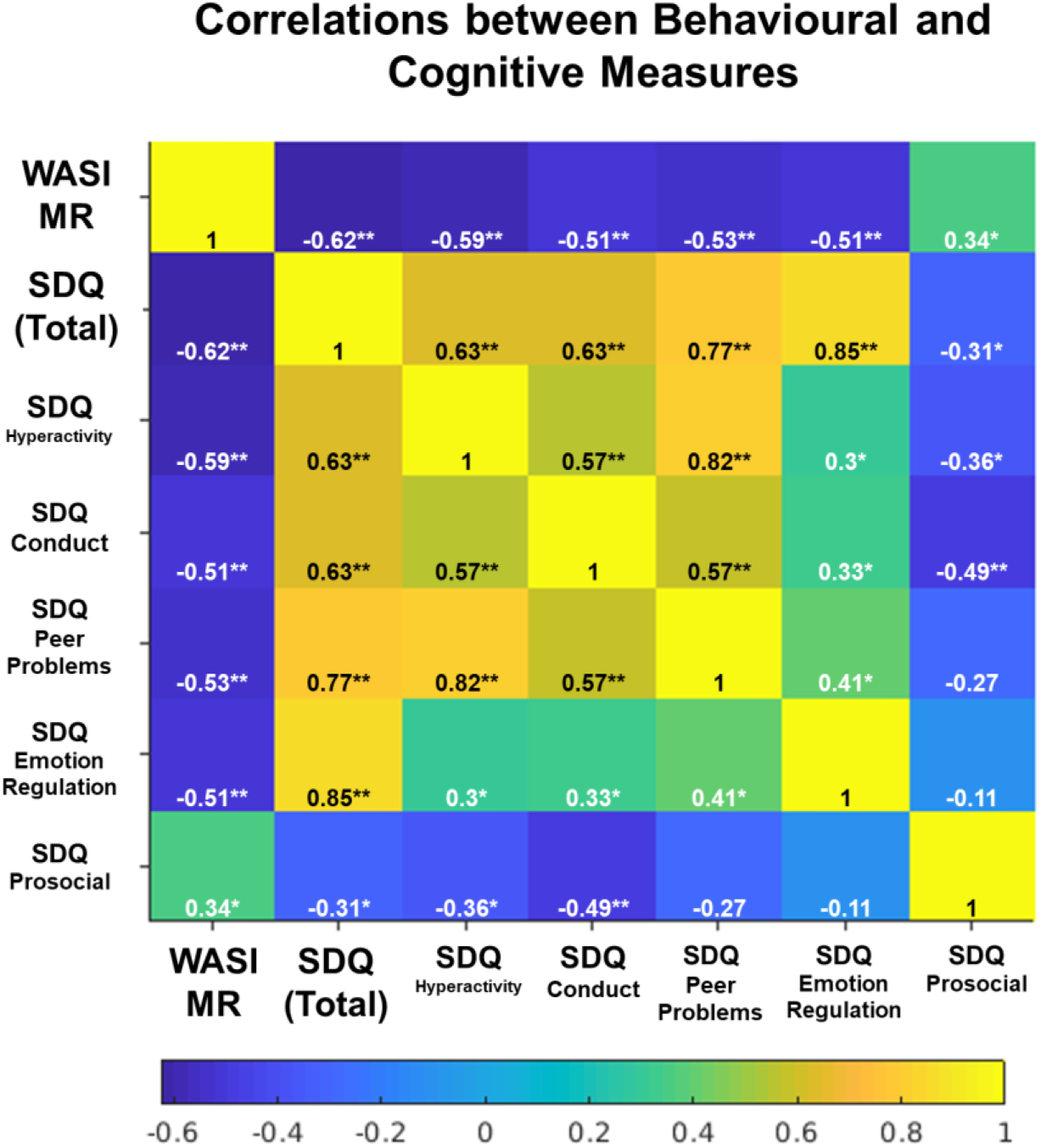
Here, we display a correlation matrix representing relationships between scores on the WASI-II MR subtest, total SDQ scores, and SDQ subscale scores. Significant relationships at p < 0.05 are marked by one asterisk, and significant relationships at p < 0.001 are marked by two asterisks.

**Table 1:** Means, standard deviations, and range values were calculated for cognitive subtest and behavioural subscale scores across our sample, which combined data from the RED study and CALM cohorts. Here, we summarise these scores, in addition to some additional sample characteristics.

| *Measure* | *Combined Sample* |
| --- | --- |
| **N** | 46 |
| **Age in Years** | 10.09 (± 1.19; range = 8-13) |
| **Gender** | 47.8% male |
| **WASI-II Matrix Reasoning** | 55.91 (± 9.70; range = 37-80) |
| **SDQ (Total)** | 7.78 (± 5.83; range = 0-10) |
| **SDQ (Hyperactivity)** | 4.11 (± 2.74; range = 0-10) |
| **SDQ (Conduct Problems)** | 1.67 (± 1.99; range = 0-8) |
| **SDQ (Peer Problems)** | 2.76 (± 2.52; range = 0-9) |
| **SDQ (Emotion Regulation Problems)** | 2.83 (± 2.9; range = 0-10) |
| **SDQ (Prosocial)** | 7.83 (± 2.58; range = 0-10) |

Correlations were performed between each of the cognitive and behavioural measures (see Figure 1). A significant negative relationship was found between WASI-II MR scores and all subscales within the SDQ (r_subscales_ < −0.59, p_subscales_ < 0.0002). Additionally, scores on the WASI-II MR were positively correlated with the SDQ Prosocial Behaviour subscale (r = 0.34, p = 0.0201).

### 3.3 Resting-state MEG acquisition

MEG data were acquired using a high-density VectorView MEG system (Elekta-Neuromag) with 102 magnetometers and 102 orthogonal pairs of planar gradiometers (306 sensors in total). Five head position indicator (HPI) coils were attached to the child’s head (one on each mastoid bone, two on the child’s forehead, and one on the top of their head) in order to monitor changes in head position throughout the recording. The positions of the HPI coils was recorded using a 3D digitizer (FASTRACK, Polhemus) in addition to over 200 additional points distributed over the scalp. Pulse was measured using an electrocardiogram electrode attached to each wrist and eye movements were recorded using horizontal and vertical electrooculograms. Data were sampled at 1Khz. Smaller children were seated on a booster seat to ensure that the tops of their heads were in contact with the scanner and that they could remain in a comfortable position for the duration of the scan.

Children were monitored by video camera throughout the scan. During the 10-minute resting-state scan, children were asked to sit as still as possible, close their eyes and let their minds wander, without falling asleep.

### 3.4 Structural MRI acquisition

Out of the 46 participants in our MEG analysis sample (following outlier exclusions), 40 participants took part in a structural MRI scan, which yielded T1-weighted images from a Siemens 3T Tim Trio system. For these images, a magnetization-prepared rapid acquisition gradient echo sequence with 1mm isometric image resolution and 2.98ms echo time was used. A natural (asymmetric) NIHPD Objective 1 scan template intended for children in pre- to mid-puberty (aged 7.5 to 13.5) was used for the 6 participants who did not undergo a T1-weighted MRI scan (Fonov et al., 2011).

### 3.5 MEG Preprocessing and Source Reconstruction

#### 3.5.1 Maxwell filtering and artefact removal

Maxwell filtering was performed using a script and repository of functions developed by Alex Anwyl-Irvine called RED Tools, which implements MNE Python’s Maxfiltering procedure (https://github.com/u01ai11/RED_Rest/tree/master/REDTools). Blinks, saccades, and pulse-related artefacts were removed by running an automated temporal independent components analysis (ICA), which applied MNE’s fastICA function to the sensor-space time-courses. Following this, we performed an additional ICA for which components were manually inspected, and any remaining ECG and EOG components were removed. Additionally, components dominated by 50Hz noise were removed to reduce electrical interference.

#### 3.5.2 Co-registration and bandpass filtering

40 participants’ MEG data from our original sample (n = 52) were co-registered to their T1-weigthed structural MRI image acquired using a 3T Siemens Tim Trio and an MPRAGE sequence. A natural (asymmetric) NIHPD Objective 1 scan template intended for children in pre- to mid-puberty (aged 7.5 to 13.5) was used for the remaining 12 participants in our original sample (6 from the RED subsample and 6 from the CALM subsample) who did not undergo a T1-weighted MRI scan. Co-registration was performed using the digitized scalp locations and fiducial markers using an iterative closest point algorithm in SPM12 (Penny et al., 2011; Wellcome Trust Centre for Neuroimaging, 2014). A forward model was fitted using a single sphere homogeneous head shape model for each subject (Mosher et al., 1999). Then, data were bandpass filtered to be between 1-30Hz in SPM12, as these slower frequencies are better for considering functional connectivity with MEG (Luckhoo et al., 2012).

#### 3.5.3 Source-localisation and parcellation

The remaining preprocessing steps were implemented using the OHBA Software Library (OSL v2.0.3; OHBA Analysis Group, 2017; https://github.com/OHBA-analysis/osl-core) and OHBA’s Hidden Markov Model Library (HMM-MAR; Vidaurre et al., 2016). First, a covariance matrix was computed across the whole time-course for each participant and was regularized to 50 dimensions using PCA rank reduction. Sensor normalisation was then performed across planar gradiometers and magnetometers. Following this, we used a linearly-constrained minimum variance beamformer to estimate whole-brain source-space activity for points in an 8mm grid (Van Veen et al., 1997). The signal-space separation algorithm reduced the dimensionality of the data, resulting in a set of estimated time-courses of brain activity for each child for 3,559 source locations across the brain (Woolrich et al., 2011). At this point in the preprocessing pipeline, we excluded 5 participants from our original sample (n = 52) who had a very high predominance of bad segments across their time-series (>60%), which was assessed using the OSL function ‘osl_detect_artefacts.m’. We excluded an additional participant on the basis of their having a poor co-registration solution. Upon visually inspecting the co-registration solution using SPM 12’s GUI, it became clear to us that scalp locations had been digitised improperly (with points placed too far from the scalp). These exclusions reduced our MEG sample from n=52 to n=46.

Following artefact-related exclusions, MEG data were further reduced down into a 38-node parcellation following the method outlined in Quinn et al. (2018). A binarised parcellation with 38 cortical regions was applied and a single time-course was estimated per node from the first principal component across voxels. This further reduced each time-course down to 38 parcels, as opposed to 3,559 voxels, and made it possible to perform additional corrections for signal leakage.

#### 3.5.4 Additional preprocessing steps

Following parcellation, further preprocessing was conducted according to OHBA’s HMM-MAR library, which recommends an additional set of preprocessing steps prior to the initialisation of the Hidden Markov Model: detrending, signal standardisation, corrections for signal leakage, and downsampling. We first completed detrending, which removes linear trends in the data for each channel separately, which was followed by a standardisation of the signal across participants’ concatenated time-courses. Next, symmetric multivariate orthogonalization was used to correct for signal leakage introduced by source reconstruction with zero temporal lag according to the methods specified in Colclough et al. (2015). Following this, the absolute signal amplitude for each source at each timepoint was estimated using the Hilbert transform. To reduce dimensionality in the data, we performed a PCA, which retained the number of dimensions necessary to explain 95% of variance in the data. Finally, MEG time-courses were downsampled to 250Hz. For a full schematic of our preprocessing procedure, please see Figure 2.

**Figure 2:**
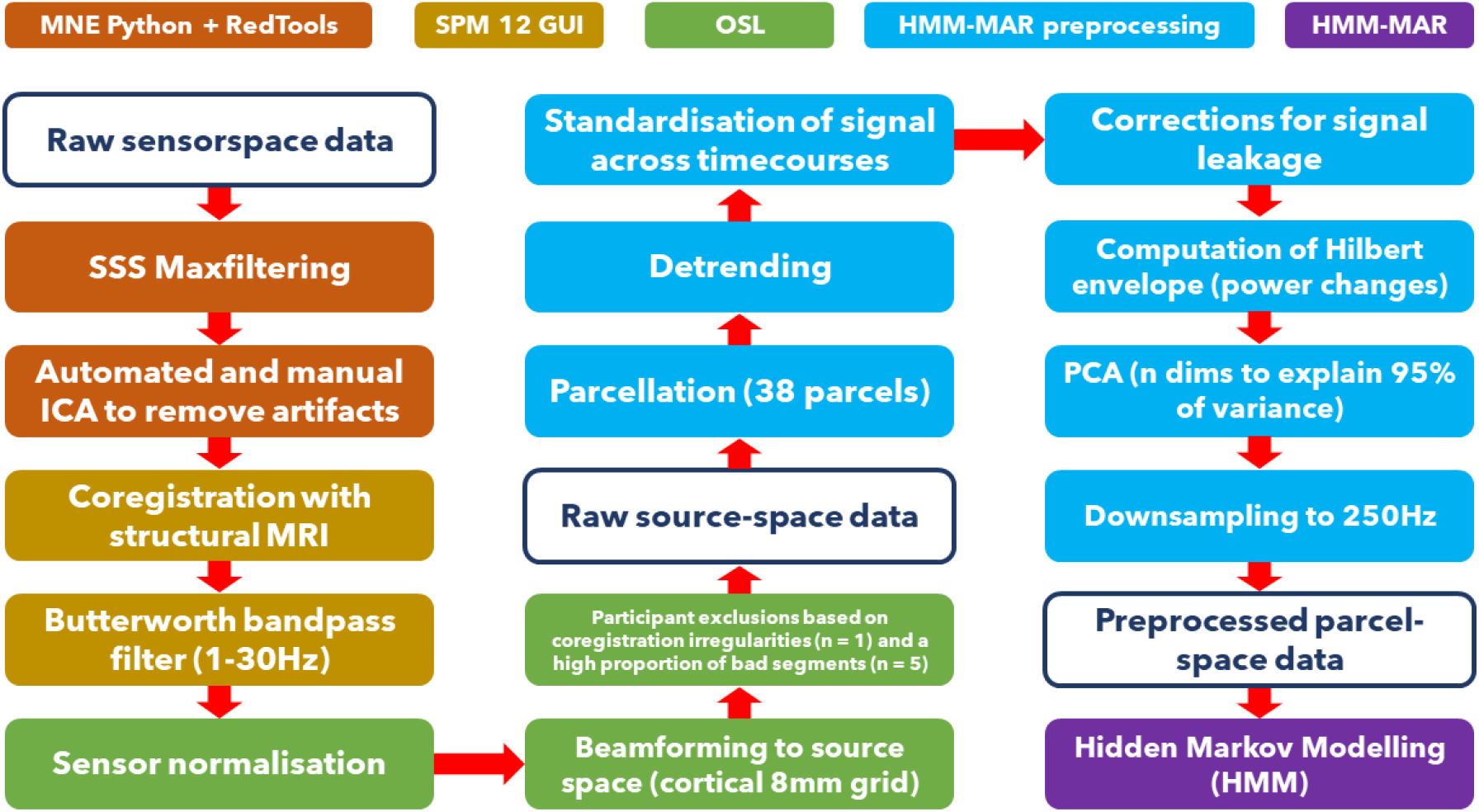
Here, we outline each step of our MEG data preprocessing pipeline, in addition to the software packages and toolboxes used to complete each step of the pipeline.

### 3.6 Hidden Markov Modelling (HMM)

In the current study, we used the HMM-MAR toolbox, developed by OHBA to infer a Hidden Markov Model from resting-state timeseries MEG data. The base code and set of functions that we adapted to suit our analyses is publicly-available on GitHub (https://github.com/OHBA-analysis/HMM-MAR). The analysis scripts for this study are also publicly-available on GitHub (https://github.com/nataliazdorovtsova/HMM_MEG).

#### 3.6.1 Model description and specifications

Hidden Markov Models (HMMs) comprise a set of unsupervised machine learning techniques that extract the spatial and temporal structure of timeseries of data by inferring discrete number of mutually-exclusive states. The HMM assumes that timeseries data, which are comprised of a set of *observed* features, can be described using a sequence of a finite, *hidden* variables (HMM states). Within a single model, HMM states are inferred on the basis that they belong to the same family of distributions, but are each parameterised differently. HMM states correspond to distinct patterns of brain activity that occur at different points across a timeseries. More formally, if we take *x_t_* to represent the time-series data and *s_t_* to represent a given state at time point *t*, we assume that

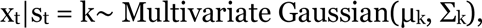

where *μ_k_* is a vector with (number of channels) elements containing the mean activation and *Σ_k_* is the (number of channels x number of channels) covariance matrix that represents the activation relationships between channels when state *k* is active. This is commonly referred to as the observational model. In this case, the observation model characterises a multivariate Gaussian distribution of each state *k* by parameters (*μ_k_*, *Σ_k_*). Although we use a multivariate Gaussian distribution to characterise states in the current study, there are a number of different methods that can be applied depending on the context-dependent theoretical goals of the researcher (e.g. Baker et al., 2014; Vidaurre et al., 2016; Vidaurre et al., 2018a; Quinn et al., 2018; Gohil et al., 2022). Here, we chose to use a multivariate Gaussian HMM with state-specific means and covariances, which can be treated as a Multivariate Autoregressive (MAR) model with an order equal to zero. This meant that the segmentation of states within our model was based on instantaneous patterns of activation and connectivity—two features of resting-state brain activity that we were interested in capturing for the purposes of this study. Previously, Vidaurre et al. (2018b) also demonstrated that multivariate Gaussian HMMs are suitable for exploring the temporal features of spontaneous transitions between large-scale resting-state brain networks.

The sequence of HMM states across a time-course is characterised by modelling the joint transition probabilities between state pairs. Put more simply, the probability *Pr* of a given state at time point *t* depends on which state was active at time point *t-1*:

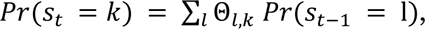

where *Θ_l,k_* refers to the transition probabilities. Within matrix *Θ*, we can further distinguish between the diagonal elements, *Θ_kk_*, which control the persistence of each state, and the off-diagonal elements, *Θ_kl_*, which refer to transitions between mutually-exclusive states. The observed data at each time point are modelled as a mixture of Gaussian distributions, with weights given by *w_tk_ = Pr(s_t_ = k)*.

In the current study, we ran a Hidden Markov Model on concatenated time-series data from all of the participants included in our dataset, which allowed us to obtain a group-level estimate of the states. Whereas the states were calculated at the group level, information pertaining to when a state is most likely to be active (the state time-course) were calculated independently for each participant.

The HMM applies an inference algorithm to estimate the parameters of each state (characterised by parameters *μ_k_* and *Σ_k_*), the probability of each state being active at each time point (*s_t_*), and the joint transition probabilities for each pair of states (*Θ_l,k_*).

#### 3.6.2 Model outputs

The *hmmmar.m* function from the HMM-MAR toolbox produces a range of outputs that can be used to estimate different features of HMM states. Below, we provide a brief description of the state properties that we used within subsequent analyses.

Following HMM inference, the temporal characteristics of each state can be quantified in terms of state fractional occupancies (the fraction of the total time spent in a state), state lifetimes (the time spent in a given state before transitioning out of that state), and interval lengths (the time it takes to re-enter a given state). Additionally, a switching rate can be calculated for each participant. Switching rates provide a measure of stability for each subject, since they represent the frequency of state switching across an individual time-course.

Using the *hmmmar.m* output *Ξ* (which holds matrices containing joint posterior probabilities of transitions between pairs of states), it is also possible to compute state transition probability matrices for each individual participant, as well as for an entire concatenated time-course. The relationships between transition probabilities and other measures of interest can be investigated in their own right, as we shall explore in the next section.

In the current study, we also used these probability matrices to derive an entropy rate estimate for each participant’s MEG timeseries data. The entropy rate measures the average uncertainty, or information, generated by a transition within a sequence. In general, the entropy rate of a sequence of random variables *(s_t_)* is defined as the limit of *H[s_0_,s_1,_…,s_n_]/n* as *n* is taken to infinity. Although this is not feasible to evaluate the entropy rate for general sequences, a sequence taken from a homogenous Markov chain posesses a compact and computationally-efficient formula for the entropy rate.

Given a transition matrix *Pr(s_t_|s_t-1_)* defining an irreducible Markov chain over a finite state space, there exists a unique invariant distribution for that chain, π, satisfying *π(s) =* ∑_𝑠′_ 𝑃𝑟(𝑠|𝑠’)𝜋(𝑠’). The entropy rate can then be computed as − ∑_𝑠′_ 𝜋(𝑠’) Pr(𝑠|𝑠^′^) log 𝑃𝑟(𝑠|𝑠’). We can understand this formula as follows: the entropy rate quantifies the average entropy of a transition within the sequence. Hence, another equivalent formula (in the case of a strongly stationary process, such as a Markov chain) is the limit of *H[s_t+1_|s_t_].* At long timescales, the convergence theorem tells us that the distribution of *s_t_* is given by invariant distribution. Hence, the entropy rate is the sum of the conditional entropies *H[s_t+1_|s_t_ = s’]* weighted by the invariant distribution 𝜋(𝑠’).

## 4 Statistical Analysis

### 4.1 HMM inference and calculation of state characteristics

HMM inference requires an a priori specification of the number of states used in the model, *k*. Free energy metrics lend some objectivity to state number selection—in theory, the ‘optimal’ number of states should be determined by the model that has the smallest free energy (measured in arbitrary units). However, it is questionable whether this practice lends itself to parsimony and theoretical coherence; the aim of the current study was to establish whether a limited collection of neural states can track differences in behaviour and cognition across our sample. Baker et al. (2014), for instance, found that free energy often increases monotonically up to *k* = 15 states, suggesting that an even higher number of states would be needed to yield a Bayes-optimal solution. A similar limitation exists for more traditional dimensionality reduction methods like Independent Components Analysis, which is driven by the goals and constraints of the research question at hand. A smaller number of prespecified components often yields canonical resting-state networks, whereas a larger number can be used to extract finer-grained distinctions between patterns of activity (Smith et al., 2011, Smitha et al., 2017).

In the current study, we trained 11 separate HMMs on our resting-state MEG dataset with prespecified states ranging from *k* = 4 to *k* = 14. Free energy metrics and information about cycles to model convergence can be found in Supplementary Table 1. After inspecting the topological features of states for each solution, we chose a HMM with *k* = 7 in order to achieve a good representation of spatially-segregated states while minimising redundancy. Varying the number of states between 4 and 14 did not appear to change the topographical features of the most prominent HMM states, which appear across solutions regardless of the addition of extra states (see Supplementary Figure 2 for plots of different model results).

### 4.2 Comparisons between measures of neural dynamics and measures of cognition and behaviour

We first used an array of General Linear Models (GLMs) to explore how participants’ switching rates, entropy rates, state fractional occupancies, and maximum fractional occupancies vary with age in order to explore developmental effects among our participants (who were 8 to 13 years old). Gender was included as a regressor in these models. Additionally, we ran a series of between-subjects t-tests to isolate any unique relationships between gender and states’ temporal properties.

We then investigated how the state measures listed above vary with measures of cognitive ability (WASI-II MR scores) and behaviour (SDQ Total and SDQ Hyperactivity, Conduct Problems, Peer Problems, Emotion Regulation Problems, and Prosocial Behaviour subscale scores). To control for potential confounds, we included gender and age as regressors in each of these models at the level of the cognitive and behavioural outcome measures.

To control for multiple comparisons, we applied a False Discovery Rate (FDR) correction with a 5% threshold in each of these analyses (see Supplementary Figures 3 and 4 for a schematic representation of the cognitive and behavioural GLMs).

## 5 Results

### 5.1 State characteristics for the seven-state HMM

A seven-state HMM revealed distinct spatial patterns of activity and variations in oscillatory amplitude (see Figure 3). Each state-map represents the mean activation profile of each parcel for the concatenated MEG dataset. State-specific increases and decreases in oscillatory amplitude are plotted as yellow/orange and cyan/blue, and represent neural activation and suppression, respectively.

**Figure 3:**
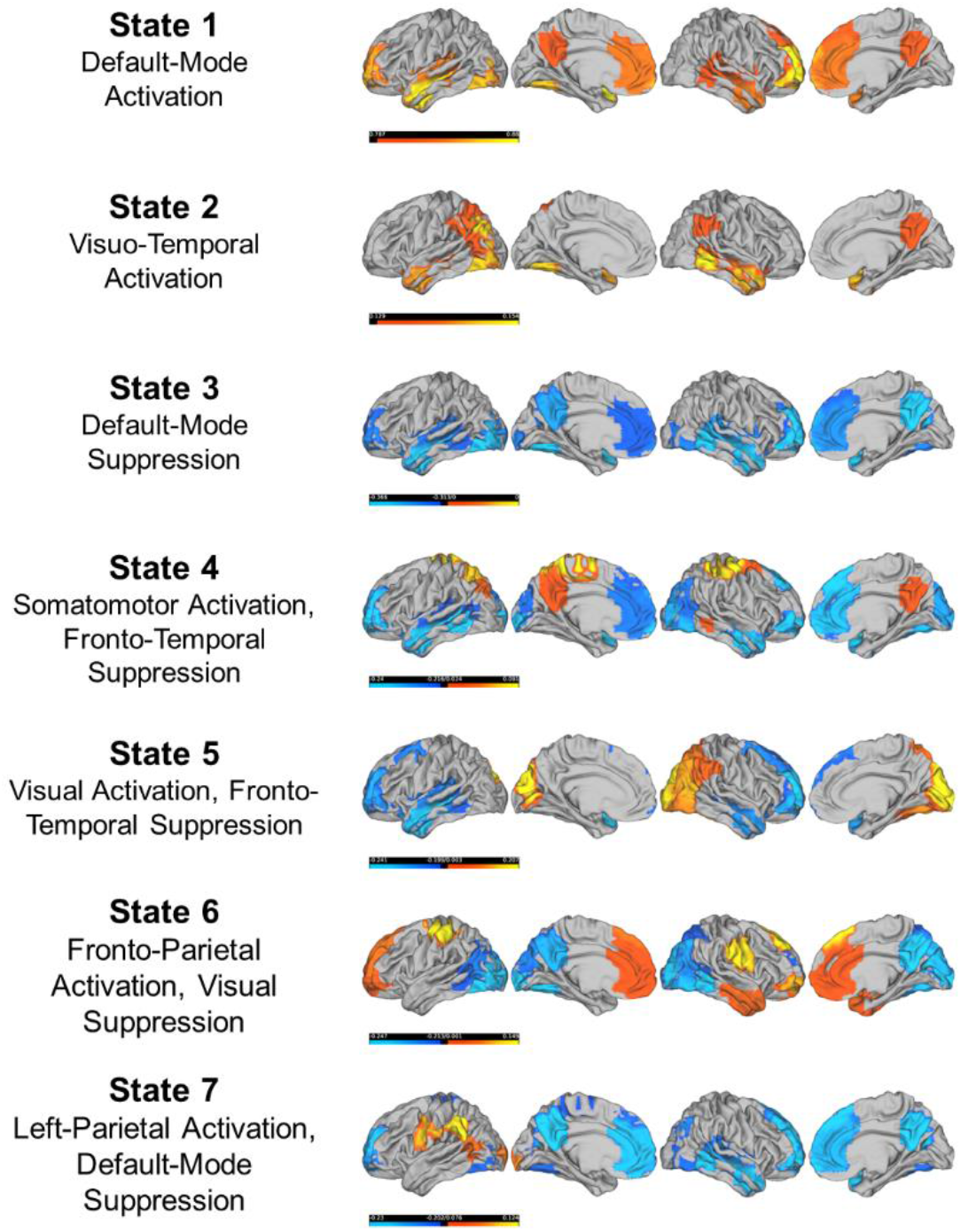
Here, we illustrate the results from our seven-state HMM. For each state, we plotted the top 20% of positive activations and bottom 20% of negative activations on a cortical surface using the HCP Workbench GUI. State labels correspond to our descriptions of the macroscopic features of cortical activation and suppression patterns.

State 1 is primarily characterised by DMN activation, and state 2 shows prominent patterns of activation in visuo-temporal regions of the cortex. States 3 and 7 both demonstrate patterns of default-mode suppression, although state 7 is also characterised by concurrent left-parietal activation. Similarly, states 4 and 5 both show fronto-temporal suppression, along with patterns of somatomotor and visual activation, respectively. State 6, meanwhile, is characterised by fronto-parietal activation and visual suppression.

Using the state time-courses, it was possible to calculate some temporal properties of each state. As illustrated in Figure 4, the temporal characteristics of the states vary considerably. Notably, state 1 has the most variable distribution in both lifetimes and intervals, which can be explained by the fact that it is the first state represented in participants’ MEG time-courses—what seems to vary, in this case, is how long participants remain in this first state that is characterised by DMN activation (see Figure 5).

**Figure 4:**
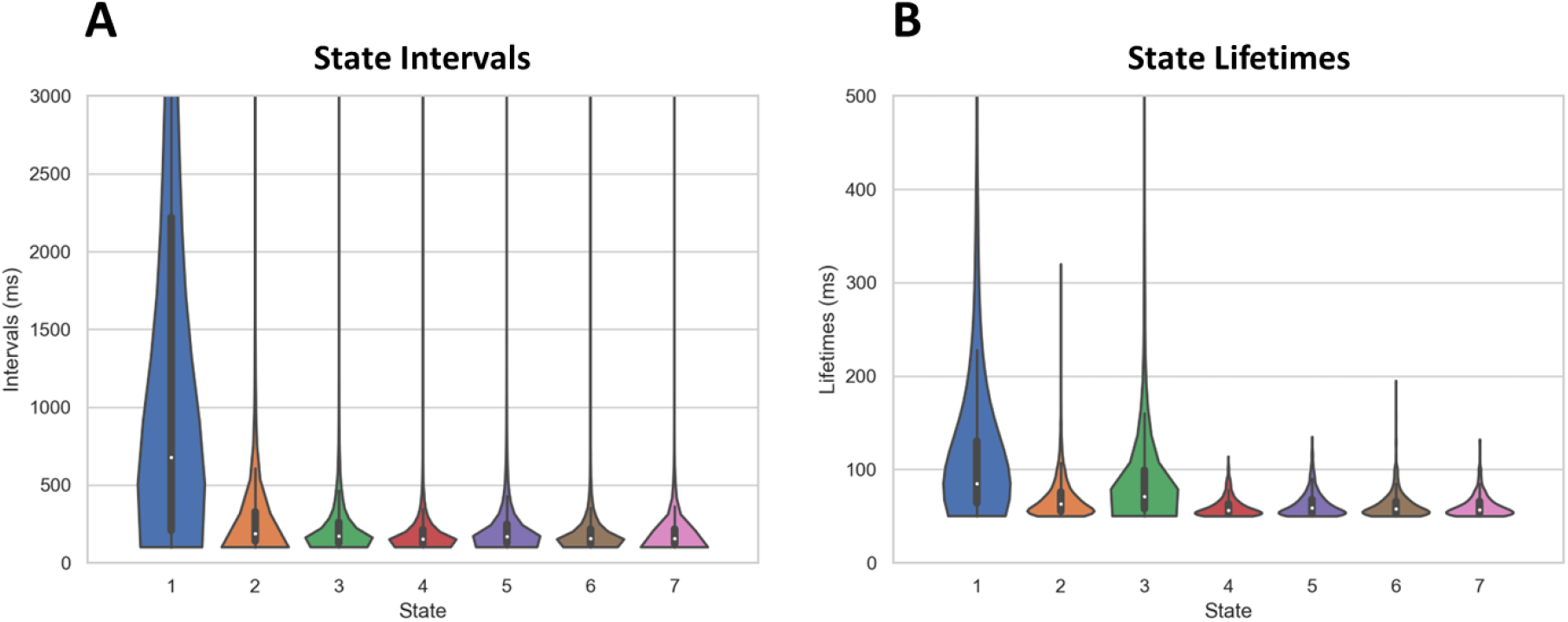
Here, we plot the intervals **(A)** and lifetimes **(B)** of the states in our k = 7 HMM. Note that state intervals and lifetimes were thresholded at 50ms, such that state appearances that were sub-50ms were not included in the calculation of these temporal metrics.

**Figure 5:**
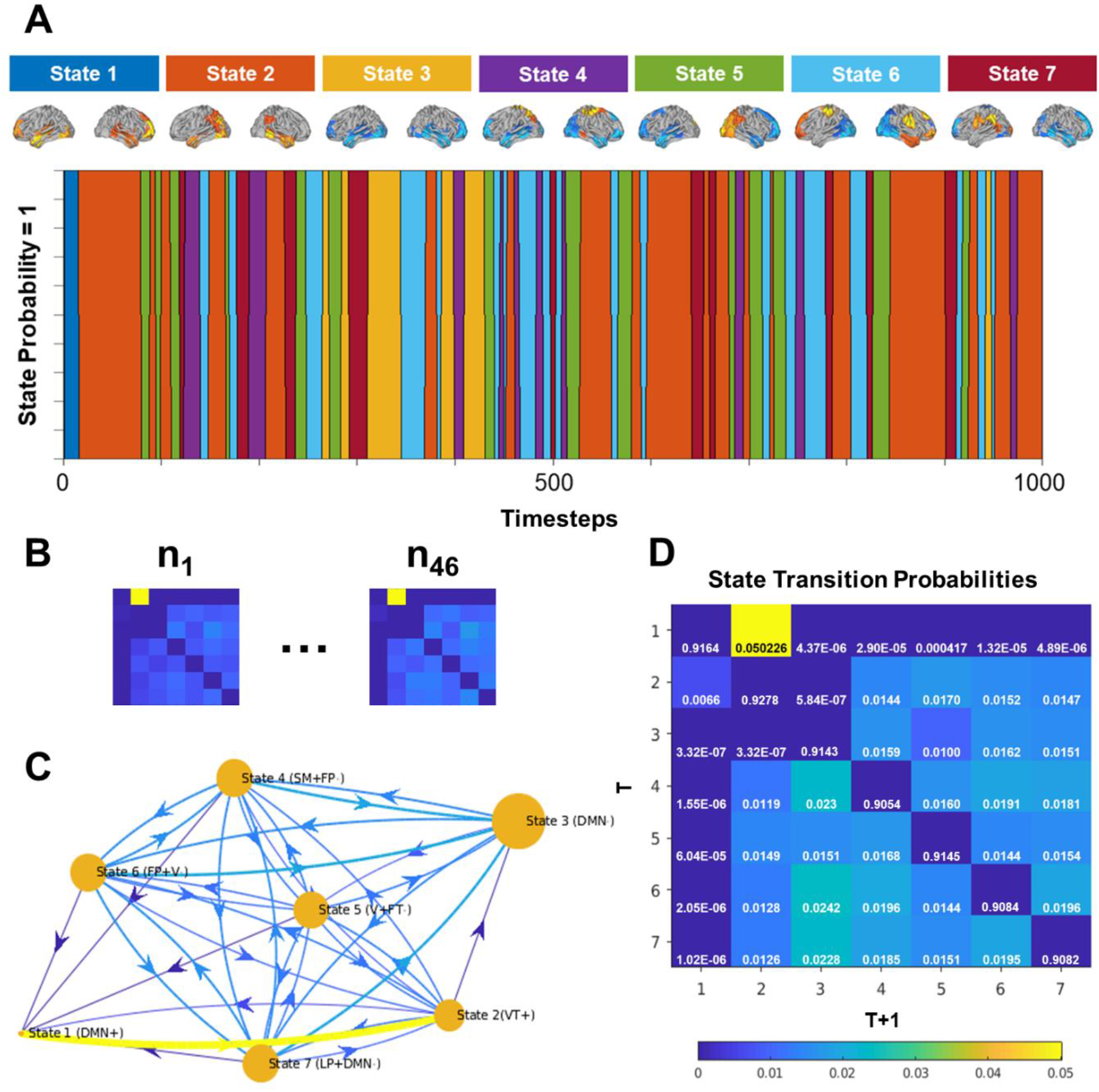
In panel **(A)**, we present the first 1000 timesteps (4 seconds sampled at 250Hz) of the Viterbi path, which represents the maximum *a posteriori* sequence of states in a HMM. Notably, state-switching is rapid, and can be detected at very short timescales. Using the joint posterior probabilities of state transitions, we computed transition matrices for each of our participants, as shown in **(B)**. Panels **(C)** and **(D)**, in which self-transitions have been intentionally excluded for the purposes of plotting state-to-state transitions, display the average transition probabilities across our entire sample (n = 46). No thresholds were applied in the generation of these plots. The node sizes in panel **(C)** reflect the fractional occupancies of each of the states.

A one-way ANOVA revealed significant differences between the states’ fractional occupancies, F_(6,315)_ = 81.42, p = 4.16 x e^-61^. Because intervals and lifetimes were calculated for each individual state, and not between participants, it was not possible to compare them in the same fashion as fractional occupancies. Nonetheless, their means and standard deviations, along with those of states’ fractional occupancies, are summarised in Table 2.

**Table 2:** Means and standard deviations (in brackets) for state intervals, lifetimes, and fractional occupancies.

| *State* | *Intervals (ms)* | *Lifetimes (ms)* | *Fractional occupancies (proportion)* |
| --- | --- | --- | --- |
| **State 1 (DMN+)** | 1925.9 (± 3344) | 129.49 (± 158.14) | 0.0177 (± 0.0145) |
| **State 2 (VT+)** | 293.9 (± 395.18) | 69.09 (± 20.90) | 0.1311 (± 0.0210) |
| **State 3 (DMN-)** | 230.38 (± 192.23) | 95.24 (± 94.50) | 0.2330 (± 0.1085) |
| **State 4 (SM+, FP-)** | 191.31 (± 143.90) | 59.75 (± 10.24) | 0.1578 (± 0.0314) |
| **State 5 (V+, FT-)** | 223.52 (± 200.73) | 63.05 (± 13.90) | 0.1524 (± 0.0229) |
| **State 6 (FP+, V-)** | 194.75 (± 149.74) | 61.99 (± 14.08) | 0.1542 (± 0.0320) |
| **State 7 (LP+, DMN-)** | 196.42 (± 170.58) | 61.20 (± 12.05) | 0.1538 (± 0.0290) |

In addition to calculating state lifetimes, intervals, and fractional occupancies, as well as generating state-maps, we also calculated state transition probabilities between and across participants. As anticipated, the most common timepoint-to-timepoint transition was the ‘self-transition’, forming the diagonal of the state transition probability matrix. State-to-state transition probabilities, which are less probable than self-transitions, are still readily-visualised when the diagonal of the matrix is zeroed-out (see Figure 5). State transition matrices with all values retained were used in later analyses.

### 5.2 Entropy-related measures of neural dynamics are related to cognitive ability, but not age, gender, or behaviour

First, we used a series of general Linear Models (GLMs) to investigate the effect of age on states’ temporal properties. Gender was included as a control regressor in these models. We did not find any significant effects of age on switching rates (t_(43)_ = 0.7221, p_adjusted_ = 0.4741), entropy rates (t_(43)_ = 0.61, p_adjusted_ = 0.5451), state fractional occupancies (t_(43)_ < |-0.5611|, p_adjusted_ = 0.9333), or maximum fractional occupancies (t_(43)_ = 0.0763, p_adjusted_ = 0.9395).

Similarly, t-tests comparing these state measures between genders did not find any significant effects for switching rates (t_(44)_ = −0.3125, p = 0.7562), entropy rates (t(44) = −0.3078, p = 0.7597), state fractional occupancies (t_(44)_ < |-0.5020|, p_adjusted_ = 0.9851), or maximum fractional occupancies (t_(44)_ = −0.1576, p = 0.8755).

Next, we investigated whether the temporal properties of states varied with six measures of behaviour (SDQ Total and SDQ Hyperactivity, Conduct Problems, Emotional Regulation Problems, Peer Problems, and Prosocial Behaviour subscales) and one measure of cognitive ability (WASI-II Matrix Reasoning). To do this, we used a series of GLMs in which age and gender were included as control regressors. No significant relationships were found between any measures of behaviour or state measures, t_(43)_ < |-2.1504|, p_adjusted_ > 0.0932 (see Supplementary Table 2 for a full summary of these non-significant results). We did, however, find significant relationships between cognitive ability and entropy rates (t_(43)_ = 2.2704, p_adjusted_ = 0.0284), and cognitive ability and switching rates (t_(43)_ = 2.1688, p_adjusted_ = 0.0358). Additionally, entropy rates and switching rates were found to be highly correlated (r = 0.9979, p < 0.00001), indicating that individual variations in entropy are almost fully explained by state switching (see Figure 6).

**Figure 6:**
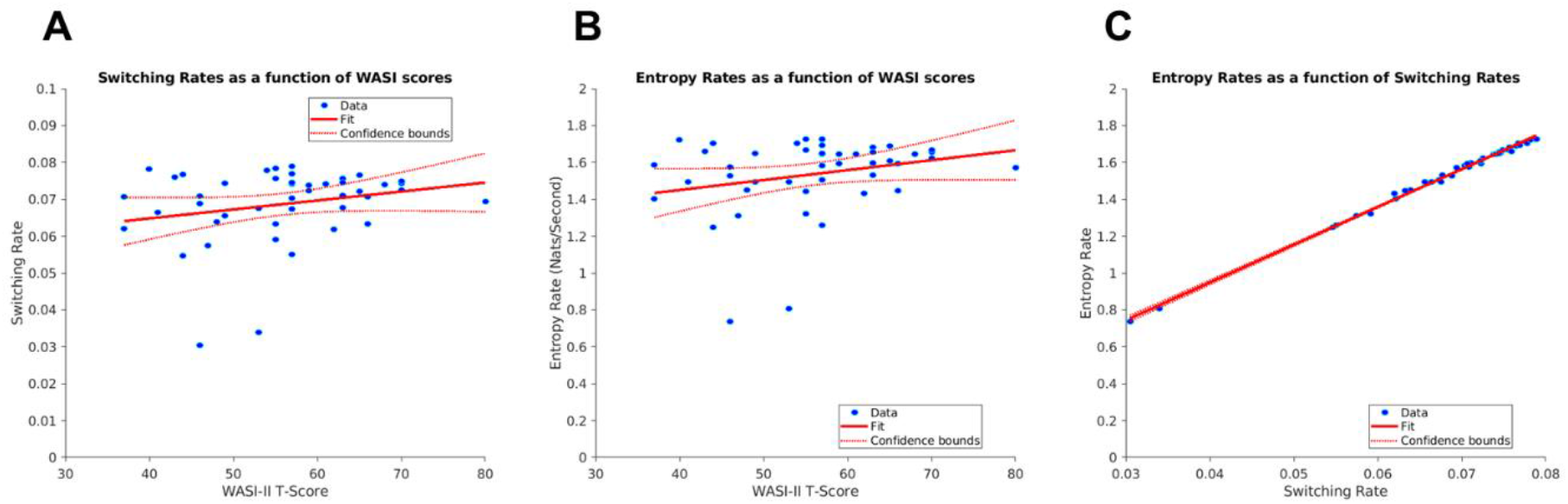
Panels **(A)** and **(B)** display the significant linear relationships between switching rates and entropy rates, respectively, and cognitive ability. Panel **(C)** shows the strong linear relationship between switching rates and entropy rates.

### 5.3 Switching is not random—state-specific fractional occupancies are related to cognitive ability

Although we found positive correlations between state switching, entropy, and cognitive ability, we wanted to investigate whether spending longer amounts of time in *specific* states could further explain the relationships between neural dynamics and cognition. Again, we used a series of GLMs in which age and gender were included as control regressors. We found significant relationships between cognitive ability and fractional occupancies in state 1 (t_(43)_ = −2.7178, p_adjusted_ = 0.0162), state 3 (t_(43)_ = −2.7693, p_adjusted_ = 0.0162), state 4 (t_(43)_ = 2.9467, p_adjusted_ = 0.0162), state 6 (t_(43)_ = 3.2673, p_adjusted_ = 0.0152), and state 7 (t_(43)_ = 2.6402, p_adjusted_ = 0.0162). An additional GLM revealed a significant relationship between cognitive ability and maximum fractional occupancies, t_(43)_ = −2.6754, p_adjusted_ = 0.0106 (see Figure 7).

**Figure 7:**
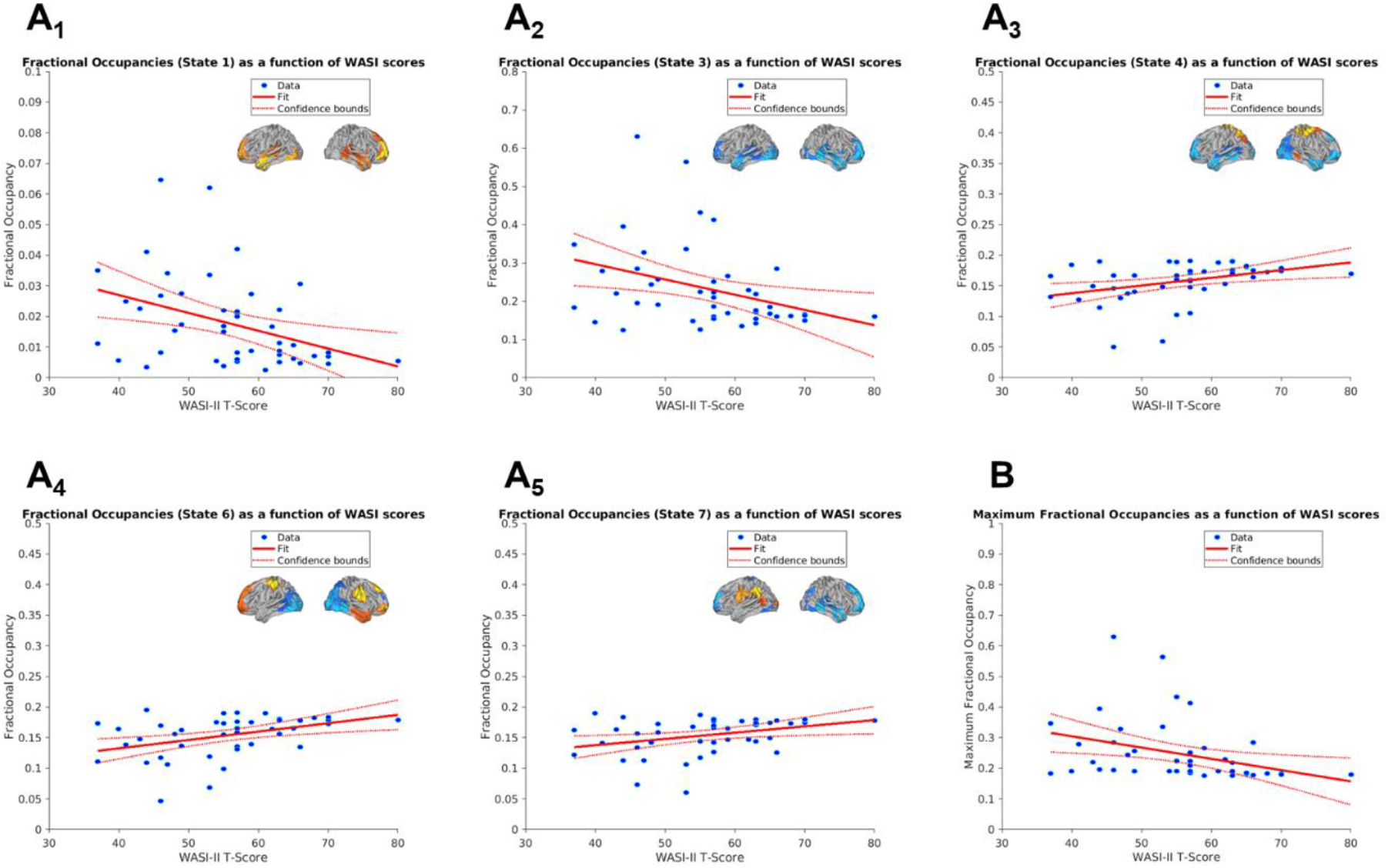
Panels **(A_1_)** to **(A_5_)** display the significant linear relationships between state fractional occupancies and cognitive ability. Panel **(B)** displays the significant negative linear relationship that was found between maximum fractional occupancies and cognitive ability.

### 5.4 Specific state transition probabilities are related to cognitive ability

Our next aim was to test whether specific state transitions were related to cognitive ability. Using the transition matrices we previously extracted for each participant, we ran a series of correlations between each cell of the 7-by-7 transition matrices and WASI-II MR scores. Although a number of significant effects at p < 0.025 were initially found (see Figure 8), indicating the presence of strong positive and negative correlations between cognitive ability and state transitions, these effects did not survive corrections for multiple comparisons at the 95% confidence interval.

**Figure 8:**
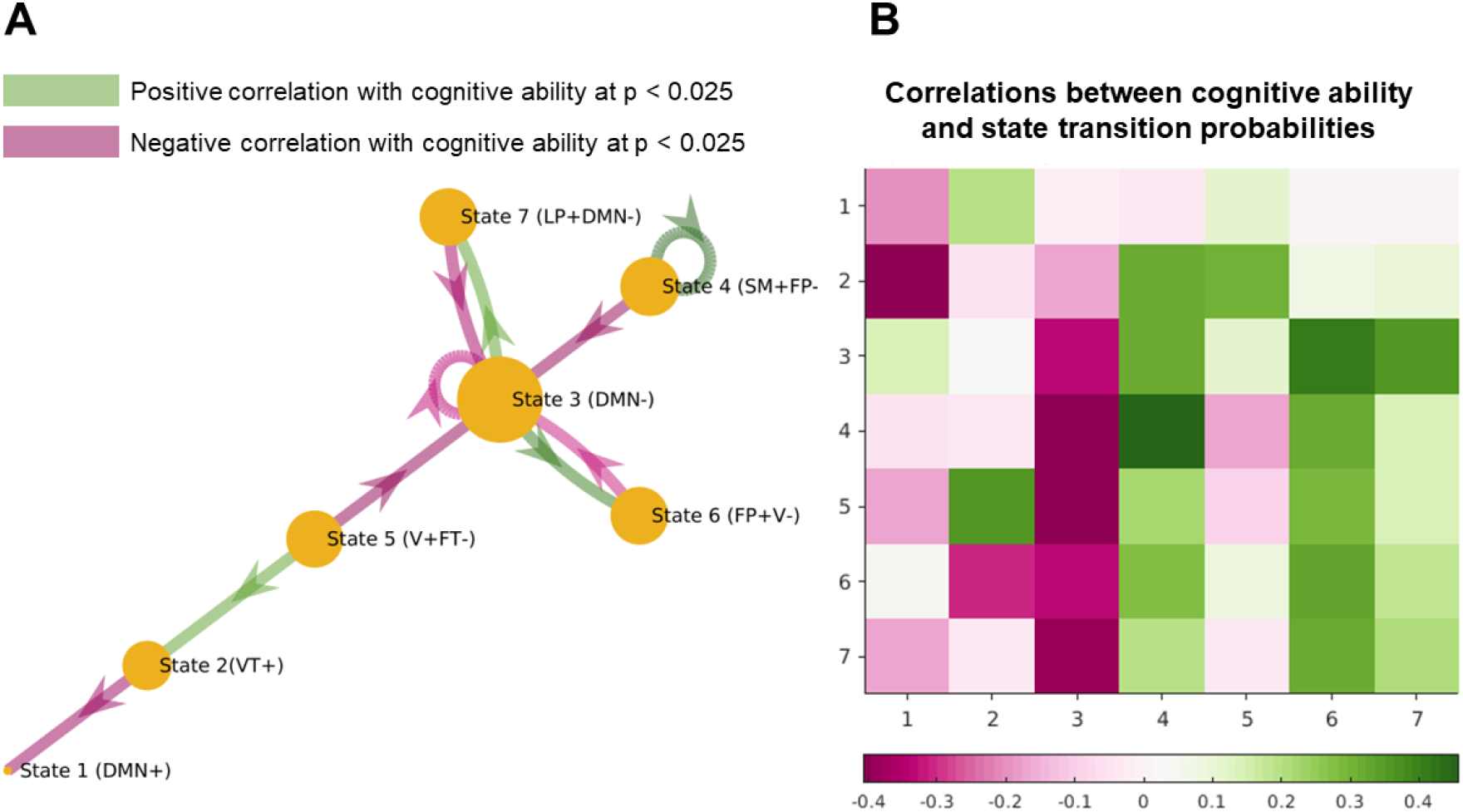
On the left, we illustrate significant relationships found between cognitive ability and transitions into specific states at p < 0.025. Node sizes represent states’ fractional occupancies, and green versus pink transition arrows represent positive and negative correlations with cognitive ability, respectively. On the right, we have plotted a heatmap representing correlations between cognitive ability and state transitions in a 7-by-7 matrix. Notably, the columns for states 3, 4 and 6 show the highest correlation coefficients, suggesting that transitions *into* these states are most strongly associated with cognitive ability.

Instead of performing correlations across 49 separate cells of the state transition matrix, we decided to investigate how transitions *into* states 1-7, represented by the columns of the matrix (which, unlike the rows, do not sum to 1), might relate to individual differences in cognitive ability. We performed 7 correlations to this effect, and found that three states were significantly correlated with cognitive ability following corrections for multiple comparisons: state 3 (r = −0.3794, p_adjusted_ = 0.0217), state 4 (r = 0.3889, p_adjusted_ = 0.0217), and state 6 (r = 0.4095, p_adjusted_ = 0.0217). We did not find any significant relationships between transitions into states 1, 2, 5, or 7 (r < |0.3154|, p_adjusted_ > 0.0573). While state 3 was negatively correlated with cognitive ability, states 4 and 6 were positively correlated with cognitive ability (see Figure 9).

**Figure 9:**
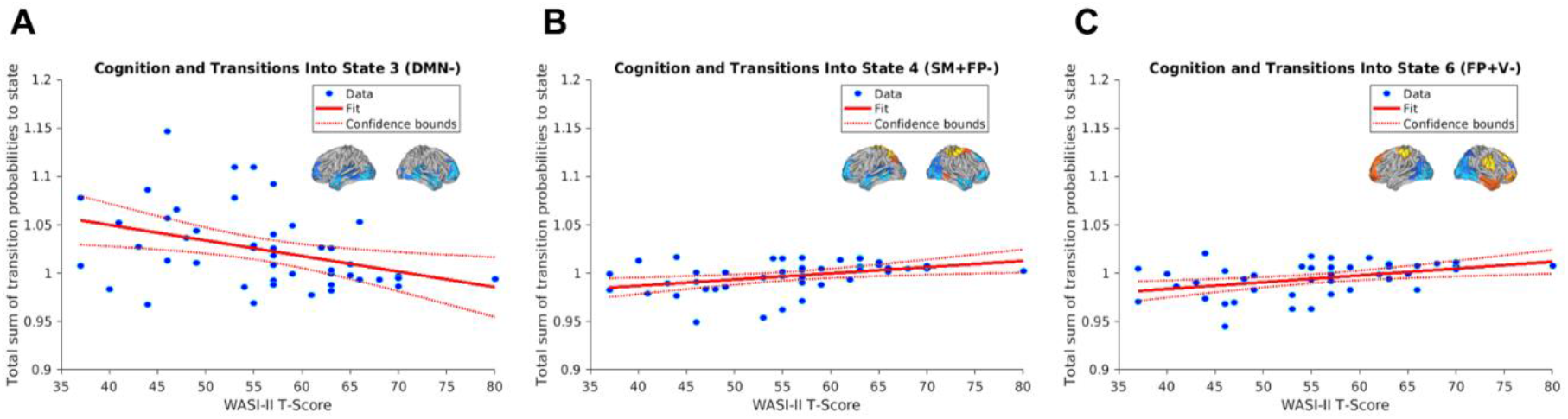
Here, we provide a finer-grained illustration of the linear relationships between transitions into specific HMM states and cognitive ability. Transitions into state 3 **(A)**, state 4 **(B)**, and state 6 **(C)** were found to be significantly correlated with cognitive ability following corrections for multiple comparisons.

## 6 Discussion

By inferring an HMM using group-concatenated MEG data from our developmental sample, we identified spatiotemporally-defined states that corresponded to well-known RSNs, including the default mode, fronto-temporal, visual, and sensorimotor networks. The spatial topographies of states in our HMM mirrored that of numerous other studies that utilised this method to extract the underlying features of resting-state MEG data, including Baker et al. (2014) and Becker et al (2020). State time-courses were characterised by a predominance of self-transitions, and that states exhibited transient (<100ms) average lifetimes. Across participants, there were individual differences in how long participants spent in each state. Cognitive ability— but not behaviour, gender, or age—was positively associated with participants’ state-switching and entropy rates. Additionally, there were state-specific relationships between cognitive ability and states’ fractional occupancies. The directionality of these relationships was preserved in analyses exploring whether transitions *into* each of the states is predictive of cognitive ability. Transitions into and time spent within DMN- heavy states was associated with lower cognitive ability, whereas the opposite was true of states with more fronto-parietal and sensory network activation profiles.

The broad alignment between our findings and those outlined in previous research suggests that an HMM approach can be applied successfully in the study of transient neural dynamics in the developing brain. Although data quality issues are known to typically affect the reliability of MEG scans in children (e.g. Wehner et al., 2008; Pang, 2011), and particularly those with high levels of behavioural difficulties (Kaiser et al., 2021), we believe our findings demonstrate that HMM inference is robust to these potential issues, provided that data are preprocessed with enough attention to noise removal and the exclusion of scans with high proportions of outliers.

### 6.1 Time-course entropy and state-switching are positively associated with cognitive ability

The switching and entropy rates of participants’ time-courses, which we found to be highly intercorrelated in our sample, were positively associated with cognitive ability (but not age, gender, or behaviour). Although state-switching and entropy were highly correlated in the current study, they have different formal definitions, and therefore have different possible interpretations. While the switching rate quantifies the extent to which a participant engages in state-to-state transitions, as opposed to self-transitions, the entropy rate can be regarded as a measure of the complexity of a participant’s time-course. Since switching rates explained the majority of the variance in participants’ entropy rates, it is reasonable to conclude that the information of MEG time-series data increased in proportion to the number of rapid and largely stochastic transitions between neural states. Had non-random and recurrent patterns of state transition sequences dominated the time-series, entropy rates would have been lower and more weakly associated with state-switching. Since the information of a time-course in our sample relies so heavily upon the flexibility and speed with which state-switching takes place, it may be useful to regard the neural entropy rate as a more complex measure that is nonetheless heavily driven by a latent switching factor.

Cortical feedback and feedforward pathways are known to rapidly control local brain network states, and this process is optimised to flexibly change the gain, precision, and synchronisation of neural activity (Zagha & McCormick, 2014). Furthermore, dynamic transitions between hidden neural states are believed to enable the flexible reconfiguration of functional circuits across the brain, thereby enabling adaptive cognitive and behavioural processes. A number of previous studies have highlighted the positive associations between state switching and cognitive ability. For instance, Taghia et al. (2018) found that task performance is strongly predicted by state-switching at rest, and that considering task-related neural dynamics only minimally improves the ability to predict task performance. Additionally, Cabral et al. (2017) found that more flexible patterns of state-switching predict better cognitive performance in older adults, and that this relationship is mediated by the tendency to transition between particular states in the neural attractor landscape. In the context of neurodevelopment, rapid switching between states is known to increase during adolescence (Medaglia et al., 2018) and accompany motor-skill acquisition (Reddy et al., 2018). Entropy, like state-switching, has also been found to increase throughout childhood development, reflecting a gradual expansion in the diversity of neural processes available to the child (Amalric & Cantlon, 2023). More broadly, brain entropy has been positively associated with general intelligence in samples with large age ranges, suggesting this brain-cognition relationship persists across the lifetime (Saxe et al., 2018; Wang, 2021; Thiele et al., 2023). Although there are many different operationalisations of brain signal complexity (e.g. Shannon entropy, multiscale entropy, Fuzzy entropy, and microstate characteristics; see Keshmiri, 2020, for review), the entropy rate of the neurophysiological HMM timeseries may also provide a useful means of exploring individual differences in the intrinsic complexity of neural activity.

### 6.2 Tendency to stay within, and transition into, certain HMM states predicts differences in cognitive ability

Although the relationship between cognitive ability and neural entropy rates was largely defined by rapid, stochastic patterns of brain switching, we wanted to investigate whether there were state-specific relationships between different neurodevelopmental characteristics and neural dynamics. While we did not find any relationships between the seven states’ fractional occupancies and age, gender, or behaviour, cognition was positively associated with time spent in states 4, 6, and 7, and negatively associated with time spent in states 1 and 3. Upon examining whether transitions into certain states also characterised these relationships, we found a similar profile of results. Transitions into states 4 and 6 had positive associations with cognition, whereas the opposite was true for state 3.

The spatial topographies of these states may provide some insight into the reason for this pattern of effects. One benefit of using MEG data to study these relationships is its high temporal resolution, which allows one to record patterns of neural activation and suppression (inhibition) that occur in very short time-windows. States 4 and 6, for instance, were primarily characterised by fronto-parietal activation *and* suppression, whereas state 3 was dominated by suppression across the default-mode network. As mentioned in the Introduction, previous research indicates that the emergence and spatiotemporal segregation of fronto-parietal networks supports executive function across development (Keller et al., 2023), whereas the overactivation and hyper-integrated spatiotemporal patterning of the DMN is associated with cognitive and behavioural difficulties (Cortese et al., 2012). The fact that both fronto-parietal activation and suppression predicts increases in cognitive ability may relate to large-scale coordination of brain activity performed by fronto-parietal networks (Marek & Dosenbach, 2018; Chen et al., 2022). Indeed, Gu et al. (2020) found that transitions to, and between, different task-positive states was positively related to performance on a cognitive task.

While DMN suppression has previously been viewed as a process that optimises for goal-directed cognition (Anticevik et al., 2012; Leonards et al., 2023), it is possible that an increased *need* to suppress DMN activity could also be viewed as a hindrance to cognitive functioning. In the current study, we found that the time spent within states corresponding to DMN activation and suppression was negatively associated with cognitive ability. The same was true of transitions from other states into the DMN- suppression state, which suggests that broad differences in DMN control may contribute to cognitive differences in childhood. To build a coherent theoretical model of the directionality of these effects, future studies of HMM states in childhood should aim to collect a wider breadth of data, and to assess how the activity of these states changes throughout development.

### 6.3 Limitations

The primary limitations of the current study are its small sample size and relatively constrained age range, which could account for why we did not find any effects of age or gender on HMM state properties. Our sample size was reduced from n=52 to n=46 due to exclusions based on data quality, which is a common difficulty within research in developmental cognitive neuroscience. While the size of our dataset was sufficient for inferring robust HMM states, it is possible that we would have been able to explore more granular neurodevelopmental effects had we had access to a larger sample. In the same vein, a sample that included a wider age range would have enabled us to investigate how neural dynamics change over time, rather than how they exist at one phase of development. We encourage future research in this area to build upon our findings with these core limitations in mind.

In this study, we also used a 38-node parcellation developed by Colclough et al. (2015) and subsequently used in other studies that applied HMM to MEG data (e.g. Colclough et al., 2017; Quinn et al., 2018). We chose this parcellation because the effective dimensionality of MEG data in source-space is approximately 60-70 (Quinn et al., 2018; Farahibozorg et al., 2018), and the number of parcels should be less than the rank of the data in order for corrections for signal leakage to work. Although using this parcellation allowed us to infer an HMM, it also reduced the spatial resolution at which we were able to observe neural effects.

## 7 Conclusion

Using a multivariate Gaussian Hidden Markov Model, we inferred a seven-state model of resting-state brain activity in a developmental sample. We found spatial and temporal differences between each of the states identified in our model. Entropy-related metrics of dynamic neural activity were positively associated with cognitive ability. Particular states provided more clarity about the nature of these relationships; DMN states were negatively associated, and fronto-parietal states were positively associated, with cognitive ability in our sample.

## Acknowledgements

The authors were supported by the Medical Research Council program grant MC-A0606-5PQ41, and D.E.A. and D.A. were supported by an Opportunity Award from the James S. McDonnell Foundation. E.J.Y. is supported by the Engineering and Physical Sciences Research Council. We would like to thank all members of the CALM and RED Teams, particularly Alexander Anwyl-Irvine, Edwin Dalmaijer, and Sophie Gibbons, for their help with recruitment, data collection, and data management. Additionally, we would like to thank all of the children and parents for their participation in the study, and the radiographers and MEG operators who support the excellent paediatric scanning at the MRC CBSU.

