## Supplement for "The entropy of resting-state neural dynamics is a marker of general cognitive ability in childhood"

**Supplementary Materials**

**Entropy-related measures of resting-state neural dynamics vary with cognitive ability in a developmental sample**

***Supplementary Figure 1****: Plots representing how cognitive, behavioural, and demographic traits were distributed across our combined sample.*

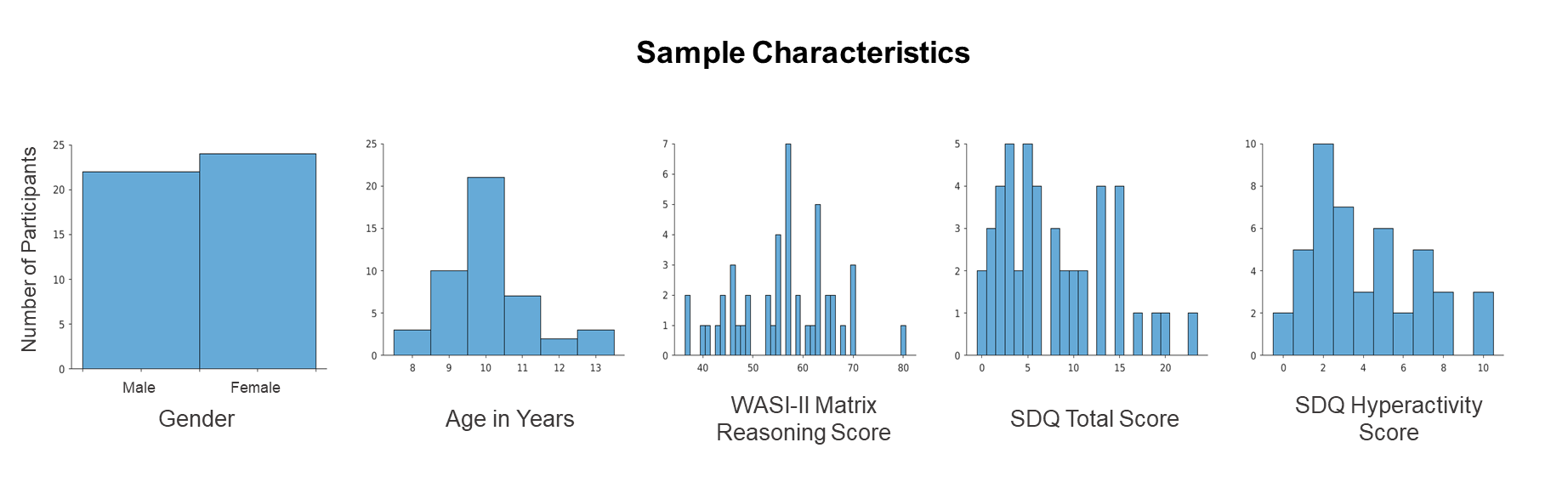

***Supplementary Table 1:*** *Output characteristics for numbers of prespecified HMM states ranging from 4-14.*

| ***Number of States*** | ***Free Energy (a.u.)*** | ***Cycles to Convergence*** |
| --- | --- | --- |
| 4 | 269430644.5 | 26 |
| 5 | 267552836 | 42 |
| 6 | 265891023.5 | 39 |
| 7 | 264251154.9 | 28 |
| 8 | 263561464.8 | 61 |
| 9 | 262547260.7 | 23 |
| 10 | 261785002.4 | 26 |
| 11 | 260785200.5 | 37 |
| 12 | 260214743.4 | 21 |
| 13 | 259412417.6 | 26 |
| 14 | 258897533.5 | 28 |

***Supplementary Figure 2:*** *Results from HMMs with prespecified numbers of states ranging from 4-14.*

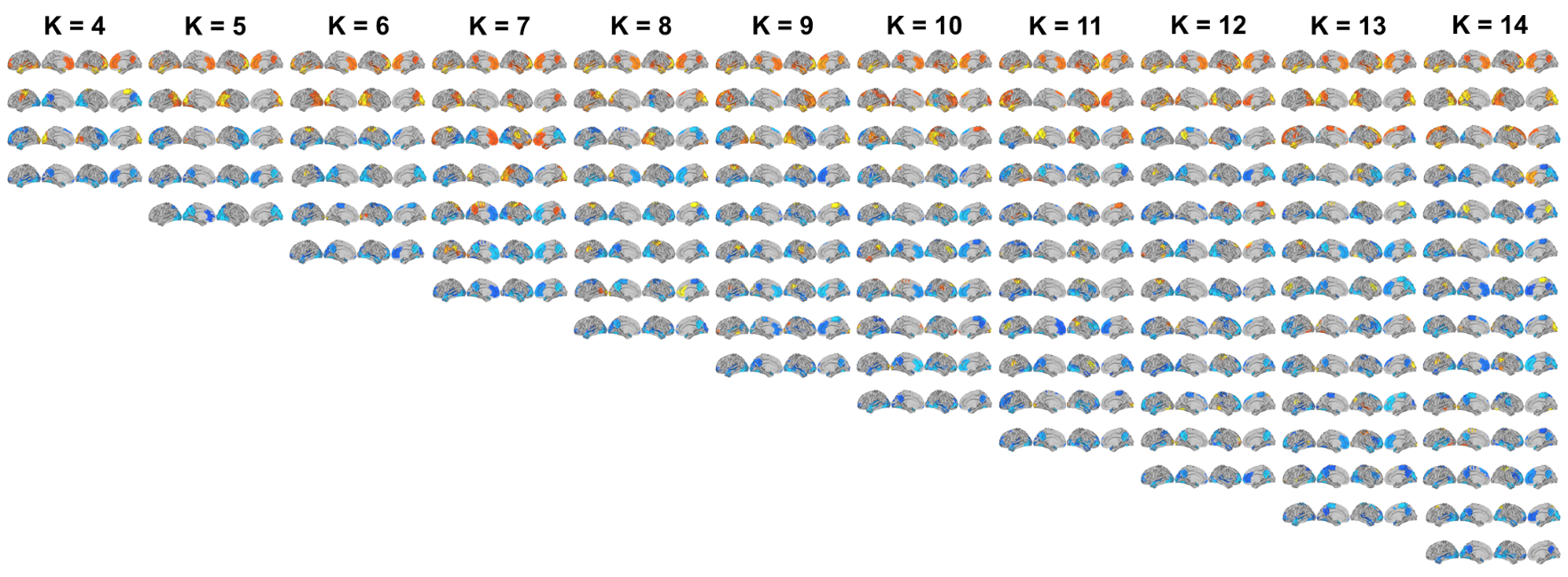

***Supplementary Figure 3:*** *Schematic representation of General Linear Model (GLM) analyses investigating the relationships between state measures and cognitive ability.*

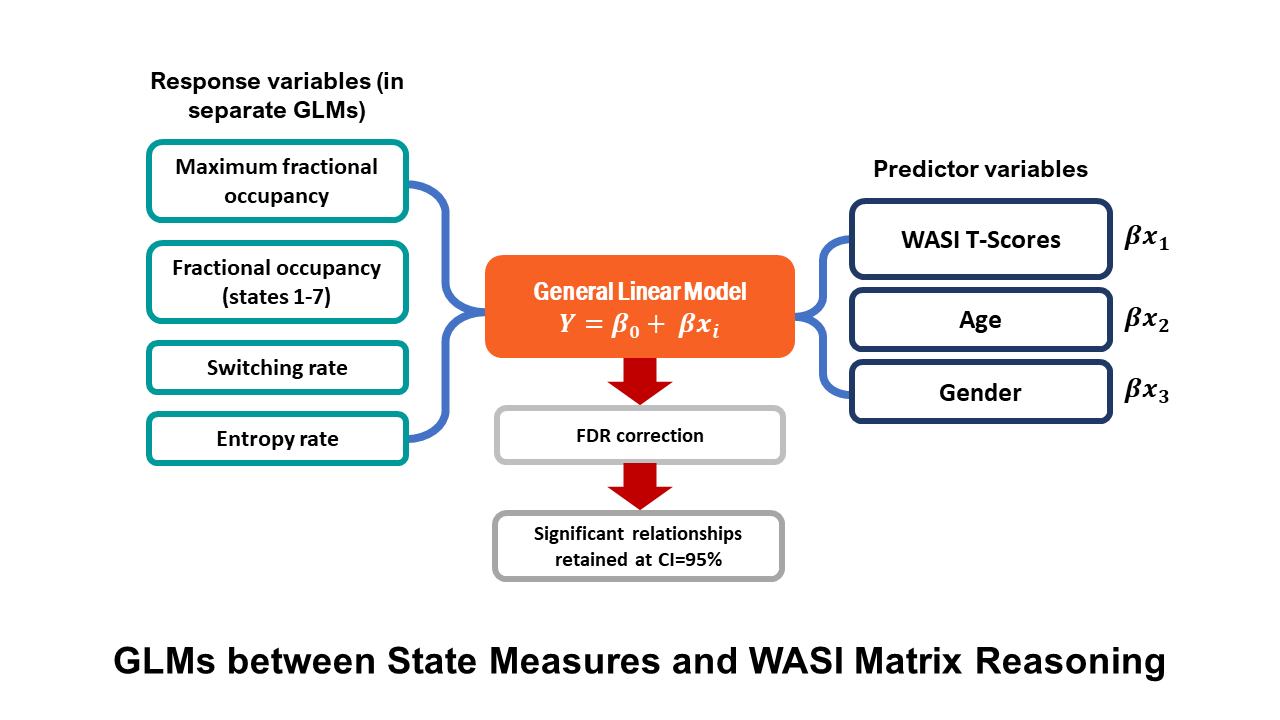

***Supplementary Figure 4:*** *Schematic representation of General Linear Model (GLM) analyses investigating the relationships between state measures and features of behaviour.*

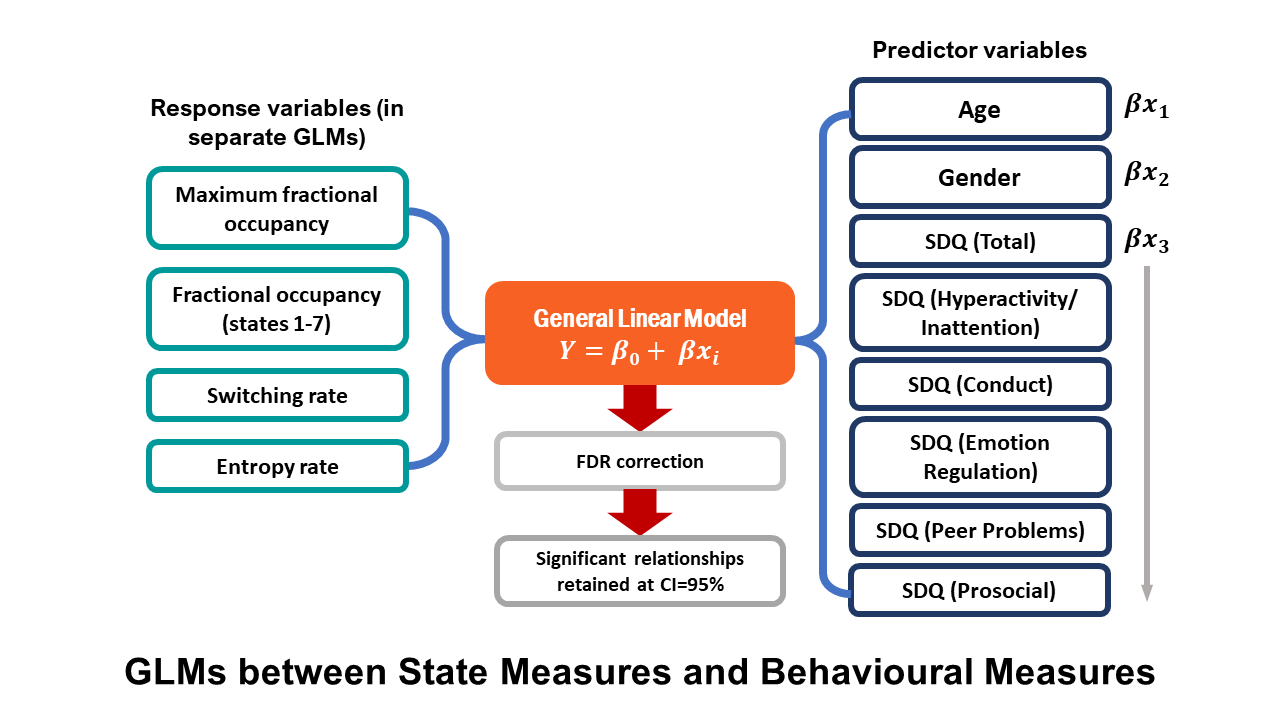

***Supplementary Table 2:*** *Results from GLMs comparing temporal state properties and measures of behaviour.*

| ***Behavioural Comparison*** | ***t-statistic*** | ***p_adjusted_*** |
| --- | --- | --- |
| **SDQ_Total_ x switching rate** | -0.7861 | 0.4362 |
| **SDQ_Total_ x entropy rate** | 0.4378 | -0.7834 |
| **SDQ_Total_ x state FOs** | <\|1.5107\| | >0.2895 |
| **SDQ_Total_ x maximum FO** | 1.1055 | o.2752 |
| **SDQ_Hyperactivity_ x switching rate** | -1.6618 | 0.1040 |
| **SDQ_Hyperactivity_ x entropy rate** | -1.6216 | 0.1124 |
| **SDQ_Hyperactivity_ x state FOs** | <\|-2.1504\| | 0.0932 |
| **SDQ_Hyperactivity_ x maximum FO** | 1.3194 | 0.1924 |
| **SDQ_Conduct_ x switching rate** | -1.3304 | 0.1906 |
| **SDQ_Conduct_ x entropy rate** | -1.3013 | 0.2002 |
| **SDQ_Conduct_ x state FOs** | <\|-2.0747\| | >0.1176 |
| **SDQ_Conduct_ x maximum FO** | 1.3350 | 0.1891 |
| **SDQ_Emotion_ x switching rate** | -0.2668 | 0.7909 |
| **SDQ_Emotion_ x entropy rate** | -0.3016 | 0.7644 |
| **SDQ_Emotion_ x state FOs** | <\|1.2939\| | >0.6885 |
| **SDQ_Emotion_ x maximum FO** | 0.8060 | 0.4248 |
| **SDQ_PeerProblems_ x switching rate** | -1.4177 | 0.1636 |
| **SDQ_PeerProblems_ x entropy rate** | -1.4162 | 0.1641 |
| **SDQ_PeerProblems_ x state FOs** | <\|2.0321\| | >0.1160 |
| **SDQ_PeerProblems_ x maximum FO** | 1.3771 | 0.1758 |
| **SDQ_Prosocial_ x switching rate** | 1.0982 | 0.2784 |
| **SDQ_Prosocial_ x entropy rate** | 0.9619 | 0.3416 |
| **SDQ_Prosocial_ x state FOs** | <\|1.4211\| | >0.3909 |
| **SDQ_Prosocial_ x maximum FO** | -0.8401 | 0.4056 |
